# Generative adversarial network (GAN) enabled on-chip contact microscopy

**DOI:** 10.1101/478982

**Authors:** Xiongchao Chen, Hao Zhang, Tingting Zhu, Yao Yao, Di Jin, Peng Fei

## Abstract

We demonstrate a deep learning based contact imaging on a CMOS chip to achieve ∼1 μm spatial resolution over a large field of view of ∼24 mm^2^. By using regular LED illumination, we acquire the single lower-resolution image of the objects placed approximate to the sensor with unit fringe magnification. For the raw contact-mode lens-free image, the pixel size of the sensor chip limits the spatial resolution. We apply a generative and adversarial network (GAN), a type of deep learning algorithm, to circumvent this limitation and effectively recover much higher resolution image of the objects, permitting sub-micron spatial resolution to be achieved across the entire sensor chip active area, which is also equivalent to the imaging field-of-view (24 mm^2^) due to unit magnification. This GAN-contact imaging approach eliminates the need of either lens or multi-frame acquisition, being very handy and cost-effective. We demonstrate the success of this approach by imaging the proliferation dynamics of cells directly cultured on the chip.

---

Lensfree on-chip imaging directly samples the light transmitted through a specimen without the use of any imaging lenses between the object and the sensor planes [1-3]. Compared to conventional lens-based microscopy, such an imaging geometry is significantly simpler and much more compact and lightweight. In addition, this geometry can decouple imaging FOV and resolution from each other, creating unique microscopes that can achieve improved resolution and FOV at the same time [4].

Contact-mode shadow imaging [5-7] and diffraction-based lensfree imaging [8-10] are the two most popular design choices for a lensfree on-chip microscope. Contact-mode imaging methods sample the transmitted light through the objects that are placed very approximate to a sensor array, effectively capturing the shadow images of the objects. Diffraction-based lensfree imaging methods, such as digital holography [11–13] and coherent diffractive imaging [14, 15] techniques, use coherent illumination to partially undo the effects of diffraction that occur between the object and the detector planes. The interference pattern formed by the scattered coherent light can be digitally processed into an image of the object. Since all these lensfree imaging modalities work under unit magnification, to mitigate pixelation-related artifacts in the digital sampling of the images, computational ways are used to increase the space-bandwidth product of the imaging devices. For instance, pixel super resolution represents a class of spatial domain techniques that use the sub-pixel shifts of the light or the sample so that a higher-resolution image with smaller effective pixel size could be created from a sequence of large FOV, low-resolution lensfree images [16–18]. Several frequency domain methods, synthetic aperture microscopy [19, 20], produce a resolution-enhanced image by stitching together a number of variably illuminated, low-resolution images in the Fourier domain. Despite offering unique imaging capabilities, these methods require special hardware setup, patterned illumination or micro-channel, and computation on multiple frames which will compromise the imaging speed. The recent advent of deep learning neural network is providing a new way to realize efficient enhancement for optical microscopy. Apart from its wide success in medical diagnosis like carcinoma detection and histopathological classification [21–24], deep learning has also been used in the super resolution for both conventional microscopy and lensfree holography [25–29].

Here we present a generative adversarial network (GAN) based contact imaging method, which is capable of resolution-enhanced lensfree imaging on a CMOS chip without the need of acquiring a plurality of frames. We simply acquire the single raw shadow image of the objects placed approximate to the sensor using ordinary incoherent illumination. A pre-trained generative and adversarial networks (GAN) can instantly recover a resolution-enhanced image of the objects in less than 1 second, achieving single-cell resolution across the entire sensor chip active area. We demonstrate the success of this GAN-enabled contact microscopy approach by imaging and analyzing the proliferation dynamics of cells directly cultured on the chip.

We used an imaging sensor (model: Aptina MT9P031) with the pitch size of 2.2 μm and active area of 5.7 x 4.28 mm for lensfree imaging. The readout part (Fig. 1b) digitalized the near-field optical signal of the cells cultured on the surface of the sensor. The obtained digital images were transferred to a PC through a USB data cable. To enable direct cell culture and onsite lensfree imaging, we carefully removed the protective glass in front of the CMOS sensor and treated the sensor with oxygen plasma for 10 minutes, to remove the micro-lens on the chip surface, thus allowing the cells to be attached approximate to the sensor surface. Then we built a water-proofing chamber onto the chip using PDMS adhesive, to accommodate the culture medium necessary for the long-term cell cultivation. The schematic and real device picture are shown in Fig. 1b. Since the multi-frame acquisition has been circumvented in our implementation, complex patterned illumination is not required in our setup. Empirically, we simply used the LED light as the illumination source. Due to the pixilation-related artifacts, the raw lens-free image usually has a large FOV while contains few high-resolution details which may be important for quantitative cell profiling. We developed a generative-and-adversarial CNN network to enable the recovery of a high-resolution lensfree image from the single low-resolution measurement. To create a well-trained GAN architecture, several high-resolution (HR) images of the cells were first obtained using a conventional high-magnification microscope (Fig. 1a, step 1). Through an image degrading model that reproduces the transfer function of the lens-free imaging process, simulated low-resolution (LR) images which perceptually analogous to the real experimental images by our lens-free device are generated (Fig. 1a, step 2)[30]. Taking HR measurements as the targets and LR simulations as the inputs, a modified generative adversarial network (GAN) iteratively learns the mapping from the low-resolution image to its corresponding high-resolution target till the goal of high-quality outputs that are close enough to the HR ground truths has been reached (Fig. 1a, step 3). It is noteworthy that this training process is required only once. Afterwards, this well-trained GAN is capable of super-resolution inference of raw lens-free measurement. Experimentally, a lensfree image of cultured cells can be quickly captured by our portable on-chip device (Fig.1b, step 1), and then inputted to the well-trained GAN for super-resolution recovery at real-time efficiency (Fig. 1b, step 2). Based on the prior knowledge learned from the training, the GAN network can quickly output a super-resolution lens-free image in less than one second. Thus, this GAN-enabled contact image keeps the large FOV from unit-magnification measurement while recovers high-resolution details which are originally decimated by the sensor pixilation.

**Fig. 1.**
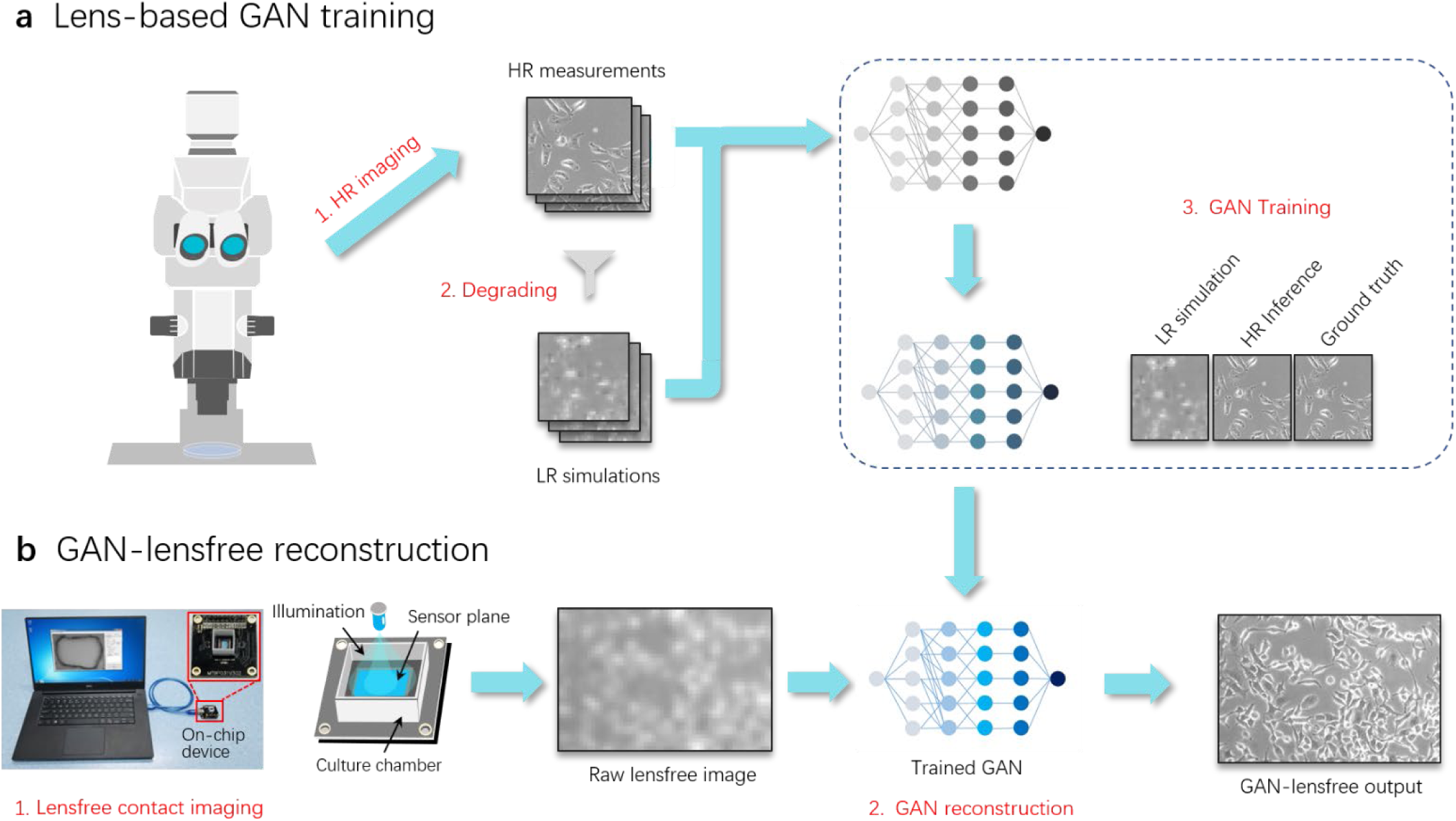
Schematic of GAN-enabled contact microscopy. (a) The procedure of lens-based GAN training including 1, Acquiring high-resolution images of the cells using a conventional microscope; 2, Generating the corresponding low-resolution simulations using our image degrading model; 3, The generative-and-adversarial training with using the low-resolution simulations as inputs and high-resolution measurements as comparative ground truths. The network iteratively learns how to optimize the HR inferences from the low-resolution simulations till the HR inferences are too similar to be discerned from the ground truths when they are compared by the discriminator. The well-trained GAN is capable of inferencing high-resolution images which are not included in the previous training dataset. (b) On-chip GAN-enabled contact imaging. A raw image with large FOV but the low spatial resolution is first captured by our lensfree on-chip imaging device. Then this single lensfree measurement is inputted to the well-trained GAN for instantly reconstructing a higher-resolution image with maintaining a large FOV of unit magnification and containing abundant cellular details which are decimated in the raw image.

The efficacy of the GAN reconstruction was first validated by the imaging test of the cells. To prove the validity of GAN, several simulated LR images (not included in the training dataset) are used as the inputs of the GAN model, of which the outputs are compared to the HR ground truths and the 4-times bicubic interpolations of the LR inputs. To further quantify the image quality improvement, we calculated the peak signal-to-noise ratio (PSNR), structural similarity (SSIM) and multi-scale structural similarity (MS-SSIM) indices between GAN results and HRs, bicubic interpolations and HRs, respectively (Tab. 1). GAN results show not only significantly better visualizations (Fig. 2) but also higher quantification scores against the interpolations, revealing its good reliability on image super resolution.

**Table 1.** Comparison of PSNR, SSIM and MS-SSIM between the Bicubic interpolation of LR inputs and GAN super-resolution outputs

|  | PSNR (dB) |  | SSIM |  | MS-SSIM |  |
| --- | --- | --- | --- | --- | --- | --- |
|  | LR | GAN | LR | GAN | LR | GAN |
| ROI 1 | 23.92 | 25.82 | 0.66 | 0.86 | 0.82 | 0.94 |
| ROI 2 | 23.19 | 25.03 | 0.65 | 0.85 | 0.82 | 0.94 |
| ROI 3 | 23.19 | 25.21 | 0.73 | 0.89 | 0.85 | 0.96 |
| ROI 4 | 23.69 | 26.06 | 0.76 | 0.91 | 0.87 | 0.96 |

**Fig. 2.**
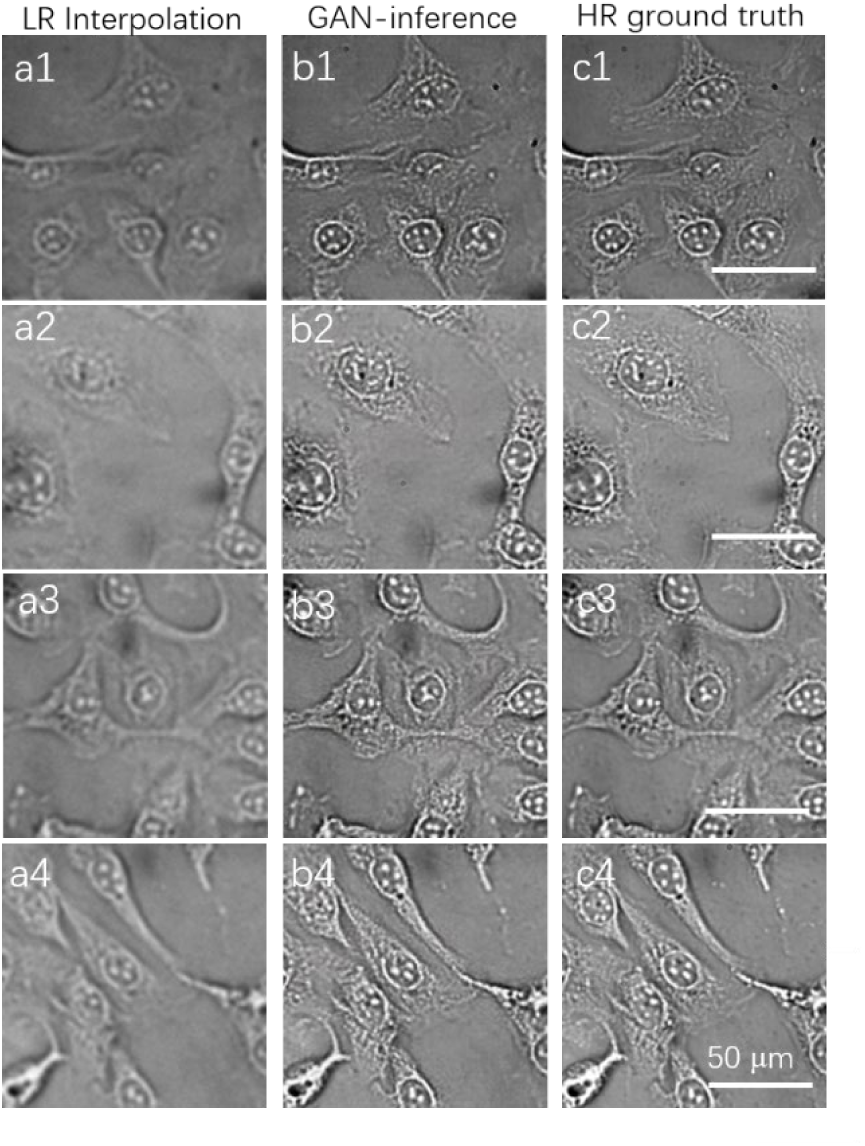
Validation of GAN super resolution. (a1) –(a4) The bicubic interpolation of the simulated LR inputs. (b1) –(b4) The GAN super-resolution results of the simulated LRs. (c1) –(c4) The corresponding HR ground truths. PSNR, SSIM and MS-SSIM indices between the resolution-enhanced (byinterpolationor GAN) versionof LR and theits corresponding HR ground truthare listed below the image.

We further demonstrate the capability of GAN-enabled contact microscopy via imaging human umbilical vein endothelial cells (HUVECs). We first performed the on-chip cell culture with complying to the standard protocol using our lensfree device. After 30-minutes sterilization of the chip by UV light, we inoculated poly-lysine-solution onto the surface of the chip to increase the cell adhesion. The remolded sensor chip was then seeded with cell medium and the whole device was put into a CO2 incubator for cell culture. After an overnight cultivation, the cells were in-situ imaged on the chip with unit magnification and then super-resolved by the well-trained GAN (Fig. 3a). The vignette high-resolution views of the cells are provided in Fig. 3b2-f2 with a reconstructed pixel size of 0.55 μm, and compared to the raw lensfree shadow image, as shown in Fig. 3b1-f1. Through the real-time GAN reconstruction (∼1s), more clear cellular profiles containing abundant features were recovered. As a result, the GAN-enabled contact imaging prototype can be considered as a compact culture/imaging device with FOV as large as the active area of the sensor chip, the same with the raw lens-free images, and maximum achieved resolution similar to that of a 20 × objective (LUCPLFLN 20X, 0.4 NA, Olympus).

**Fig. 3.**
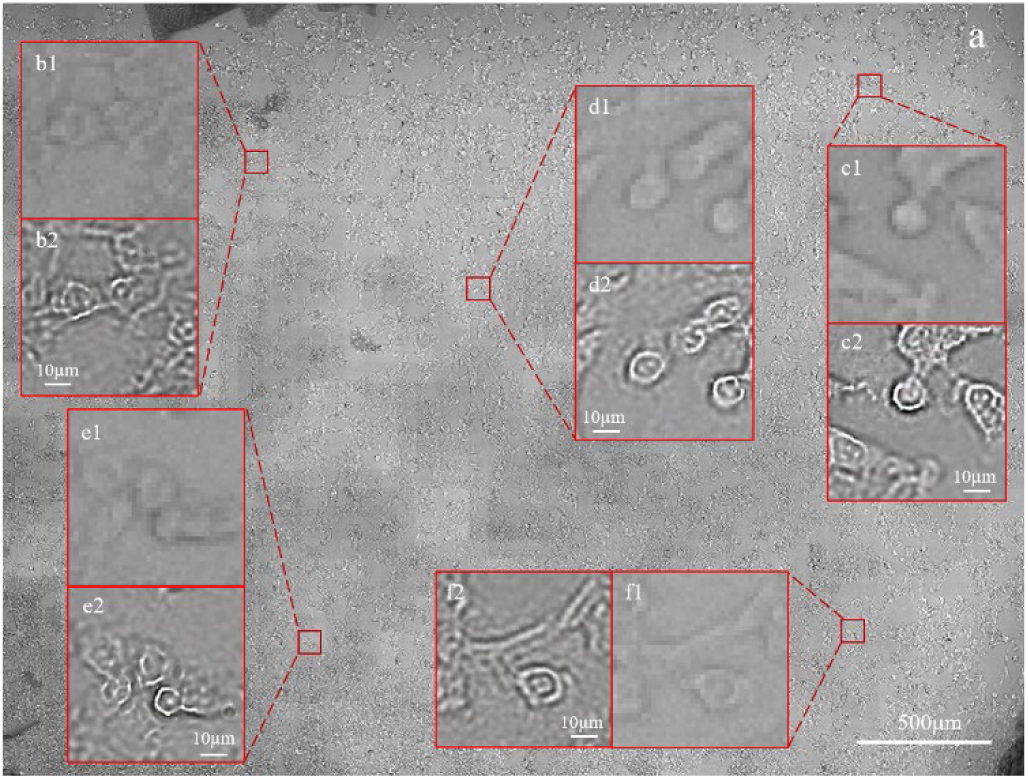
Large-FOV and high-throughput GAN-enabled contact imaging. (a) GAN-reconstructed lensfree image of HUVEC after 24 hours’ on-chip culture. (b1) -(f1) Images from the raw lensfree shadow image, for comparison. (b2) -(f2) Vignette high-resolution views of the image in(a).

**Fig. 4.**
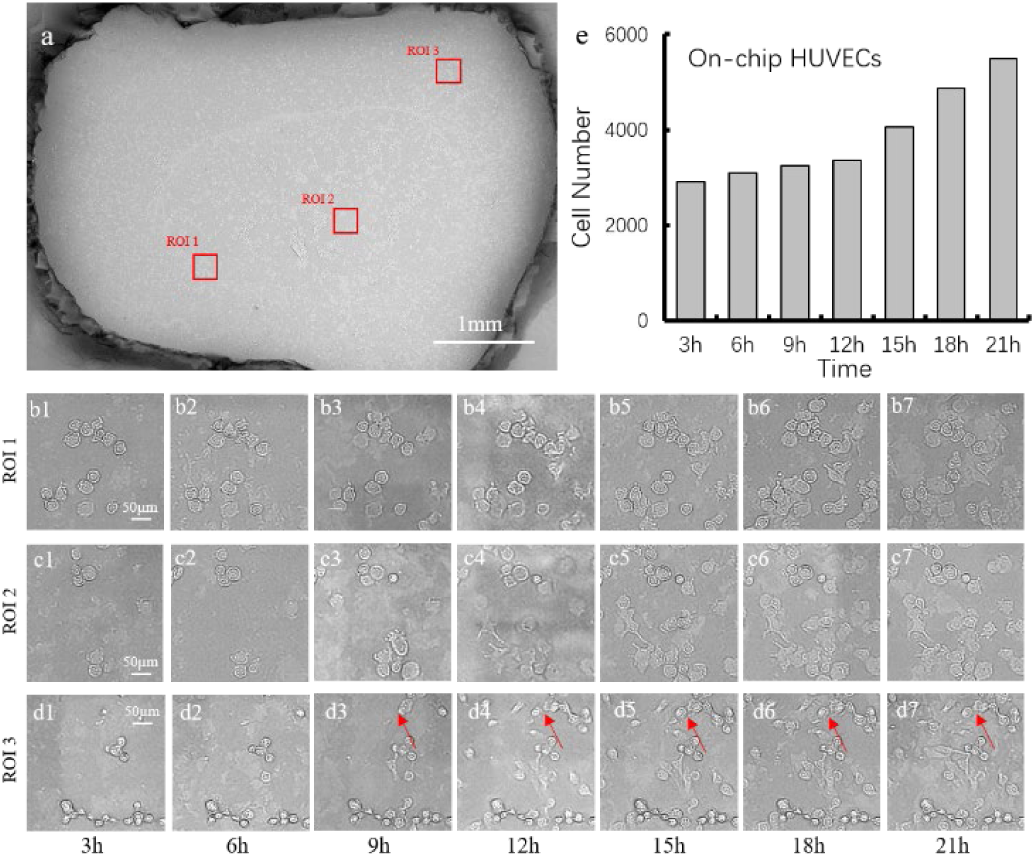
The proliferation dynamics of cells on the chip. (a) A GAN-enabled contact image after 6 hours’ cell culture. (b) –(d) Three regions of interest (ROIs) selected to reveal the time-varying dynamics of cell proliferation.(e) The total number of the on-chip cultured cells counted.

Besides the end-point detection, we also imaged the dynamics of living cells on chip over time. The cultured cells were imaged on chip every 3 hours till the endpoint of 21 hours. At each time point, we obtained a large-FOV, high-resolution GAN image for subsequent cell counting. For each GAN reconstruction, a number of regions of interest (ROIs) were selected to reveal the time-varying dynamics of cell proliferation, as shown in Fig. 3b, c, d. Benefiting from the improved resolution, the cells indicated by the red arrows in the Fig. 3d3 to d7 were observed to complete a cycle of division. The population change of the cells on the whole chip was also calculated over the entire culture period, as shown in Fig. 3e. It shows that at the first 9 hours of culture, the number of cells increased from ∼2907 to ∼3239, which is slower than the increase at later period (from 3366 to 5493) probably due to the necessary process of cell adhesion at the initial stage.

We have demonstrated a deep learning-enabled contact imaging method, which can computationally improve the resolution of raw lensfree images, and thus increase the optical throughput for large-scale on-chip imaging. Once the Generative-and-adversarial network being sufficiently trained using lens-based dataset, it is capable of reconstructing a large-FOV, super-resolution image based on single pixelated lensfree measurement, in real time. GAN-enabled contact imaging extends the space bandwidth product of conventional lensfree device without the cost of acquiring multiple frames using extra patterned illumination. As a demonstration, we proved it has a chip-size FOV (∼24 mm2) to accommodate large-scale cell culture and improved single-cell resolution (∼1 μm) for quantitative analysis. The performance could be further improved by using better sensor chip. In addition, it is easily reproducible and highly cost-effective for resource-limited environments. Due to the compact form factor, the device can be easily integrated into an incubator to conduct long-term, on-site observation of living cells, giving further insight into the time-course study of cell dynamics. These advancements render GAN-enabled contact imaging method a potential tool for a variety of biomedical assays, such as invitro drug test and stem cell differentiation & migration, in which cellular profiling at large-scale is highly desired.

## Funding

National Key R&D program of China (P.F., 2017YFA0700500), The National Natural Science Foundation of China (21874052), Research Program of Shenzhen (P.F., JCYJ20160429182424047), 1000 Youth Talents Plan of China (P.F.).

## Acknowledgements

The authors acknowledge the selfless sharing of the GAN source codes from Hao Dong. The authors thank Wenbin Jiang and Yang Ma for their assistance with GPU-based computation.

